# Homeoprotein ENGRAILED-1 promotes motoneuron survival and motor functions

**DOI:** 10.1101/734020

**Authors:** Stephanie E. Vargas Abonce, Mélanie Lebœuf, Kenneth L. Moya, Alain Prochiantz

## Abstract

Motoneuron degeneration leads to skeletal muscle denervation and impaired motor functions, yet the signals involved remain poorly understood. We find that extracellular ENGRAILED-1, a homeoprotein expressed in spinal cord V1 interneurons that synapse on α-motoneurons, has non-cell autonomous activity. Mice heterozygote for *Engrailed-1* develop muscle weakness, abnormal spinal reflex and partial neuromuscular junction denervation. A single intrathecal injection of ENGRAILED-1 restores innervation, limb strength, extensor reflex and prevents lumbar α-motoneuron death for several months. The autophagy gene *p62*, which was found to network with *Engrailed-1* and amyotrophic lateral sclerosis genes, is misregulated in *Engrailed-1* heterozygote mice α-motoneurons and is rescued following ENGRAILED-1 injection. These results identify ENGRAILED-1 as an α-motoneuron trophic factor with long-lasting protective activity.

## INTRODUCTION

Discovered on the basis of their developmental functions, homeoprotein (HP) transcription factors remain expressed in the adult where they participate in physiological homeostasis(*1*). HP functions involve cell autonomous and non-cell autonomous activities, a consequence of their ability to transfer between cells (*2*). Examples of *in vivo* non-cell autonomous activities include the regulation of cerebral cortex plasticity by OTX2 (*3*) the control of Cajal-Retzius cell migration by PAX6 (*4*) and the axon guidance activities of ENGRAILED and VAX-1 (*5*, *6*).

In *Engrailed-1* heterozygote (*En1-Het*) mice, mesencephalic dopaminergic (mDA) neurons in the Substantia Nigra (SNpc) die progressively starting at post-natal week 6 (*7*). Infusion or injection of ENGRAILED-1 (EN1) or EN2 protein in the SNpc, and its internalization by mDA neurons, stops their degeneration in several Parkinson Disease (PD) mouse and non-human primate models (*8*, *9*). A single EN1 injection in mice is active for several weeks, reflecting that EN1, not only regulates transcription and translation, but also works at an epigenetic level (*10*, *11*).

*En1* is expressed in developing and early postnatal spinal cord V1 inhibitory interneurons that form inhibitory synapses on large α-motoneurons (αMNs) and regulate alternating flexor-extensor activity contributing to the excitatory/inhibitory balance important for αMN survival (*12*, *13*). In light of EN1, EN2 and OTX2 neuronal pro-survival activities (*2*), we investigated whether EN1 from V1 interneurons impact on αMN physiology and survival. We demonstrate that *En1* expression is maintained in adulthood, that *En1-Het* mice show early muscle weakness with partial denervation of the neuromuscular junction (NMJ) and a loss of αMNs. These deficits increase after 2 to 3 months of age and are, in part, reproduced by blocking EN1 non-cell autonomous activity. A single intrathecal injection of human recombinant EN1 (hEN1) at 3 months restores muscle strength, normalizes spinal reflex and NMJ innervation and promotes αMN survival. This restorative activity lasts for at least 3 months.

## RESULTS

### Neutralizing extracellular EN1 in the adult induces motor deficits

*Engrailed-1* is expressed in a large population of neurons dorsal to the main population of large ChAT-stained MNs and in the ventral Calbindin-1 positive Renshaw cells at 4.5 months (Fig. 1A). *This* expression is identical at 4.5 and 16 months and reduced by half in the *En1* heterozygote. Thus, large ChAT-expressing MNs are surrounded by *En1*-expressing cells.

**Figure 1.**
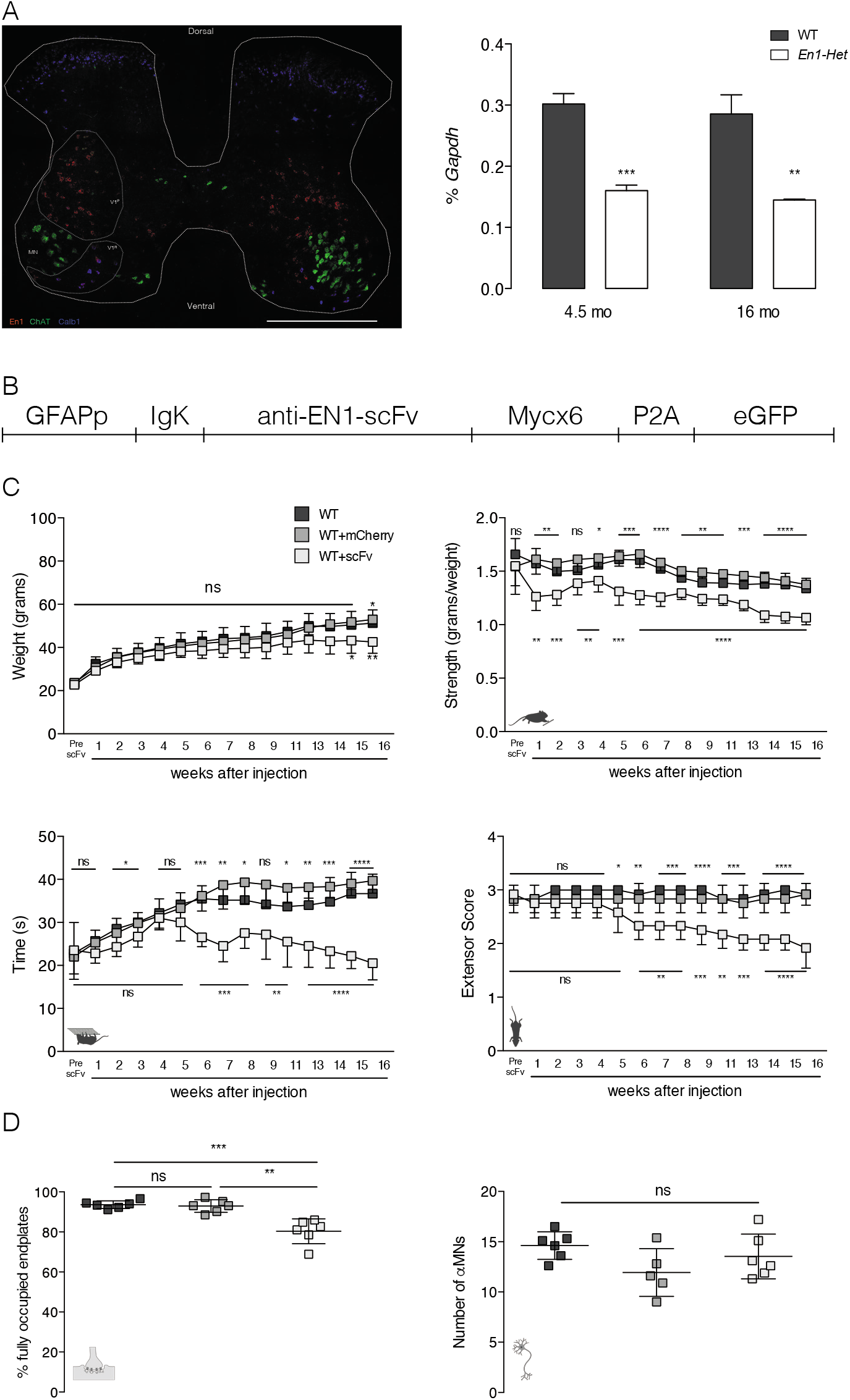
Extracellular EN1 neutralization decreases muscular strength and modifies synapse morphology. **A**. *In situ* hybridization showing *Engrailed-1* (*En1*), *Choline acetyl transferase* (*ChAT*) and *Calbindin-1* (*Calb1*) gene expression in the lumbar spinal cord. *En1* is expressed in V1 interneurons, dorsal (V1^P^) and ventral (V1^R^) to the main motoneuron pool (MN). Ventral interneurons correspond to Renshaw cells. Scale bar: 500 μm. RT-qPCR of RNA from the lumbar enlargement of 4.5- and 16-month-old shows stable *En1* expression in WT at both ages and reduced expression in heterozygous mice. *En1* probe specificity was verified on *En1*-expressing midbrain structures (supplemental Fig.2a) and EN1-het mice show less EN1 expression in spinal interneurons (Fig 2b). **B**. AAV8-encoded construct containing glial fibrillary acidic protein (GFAP) promoter, Immunoglobulin K (IgK) signal peptide for secretion, anti-ENGRAILED single-chain antibody (AntiEN1-scFv), 6 myc tags (6xMyc), skipping P2A peptide and enhanced Green Fluorescent Protein (eGFP). **C**. scFv expression has no effect on weight gain before week 15 (top left). Forelimb grip strength is reduced after week 1 (top right) and time on the grid (bottom left) and extensor reflex (bottom right) at week 6. Statistics in the top and bottom graphs compare scFv-exressing mice to WT and mCherry-expressing mice, respectively. **D**. Extracellular EN1 neutralization (16 weeks post-infection) decreases the percentage of fully occupied endplates (left) but does not affect the number of αMNs (right). Unpaired T-test with equal SD (*p<0.05; **p<0.005; ***p<0.0005; ****p<0.0001). n=6 for all groups.

ENGRAILED1/2 is a transduction protein that directly signals between cells (*2*). Neutralizing extracellular EN1/2 *in vivo* modifies the maturation of the anterior cross vein in the fly wing disk and the navigation of retinal ganglion cell growth cones in frog and chick tectum (*5*, *14*). In both cases, extracellular EN1/2 was neutralized by expression of a secreted single-chain antibody (anti-EN1/2 scFv). We neutralized extracellular EN1 in the spinal cord, using an AAV8 encoding the myc-tagged anti-EN1/2 scFv under the control of the astrocyte-specific Glial Fibrillary Acidic Protein (GFAP) promoter (Fig. 1B) injected intrathecally at the lumbar level in 1-month-old WT mice.

Figure 1C shows the effects of the anti-EN1/2 scFv on weight, forepaw grip strength, holding onto an inverted grid and hindlimb extensor reflex. A minor weight loss appeared late after injection, whereas muscle weakness and abnormal hindlimb reflex appeared early and deteriorated with time. Neurofilament and synaptic vesicle protein 2A antibodies were used to visualize the innervation of the motor endplates characterized by alpha-bungarotoxin binding to acetylcholine receptor (AChR) clusters (Supplemetal Fig. 1A). Full innervation was reduced 16 weeks following scFv injection (Fig. 1D), but not the number of αMNs identified by size and ChAT expression (Supplemental Fig. 1B-D).

### Phenotypic characterization of the *En1*-*Het* mutant mouse

In absence of weight differences, *En1-Het* animals showed progressive loss of strength compared to their WT littermates (Fig. 2A-D). Forepaw grip strength was reduced at 3 months and persisted through 15.5 months of age. In the inverted grid test, *En1-Het* mice held on significantly less time at 2 months than WT mice with further deterioration over time. Finally, the hindlimb extensor reflex deteriorated between 1 and 9 months. Taken together, *En1-Het* mice show a rapid period of strength loss between 1 and 4.5 months.

**Figure 2.**
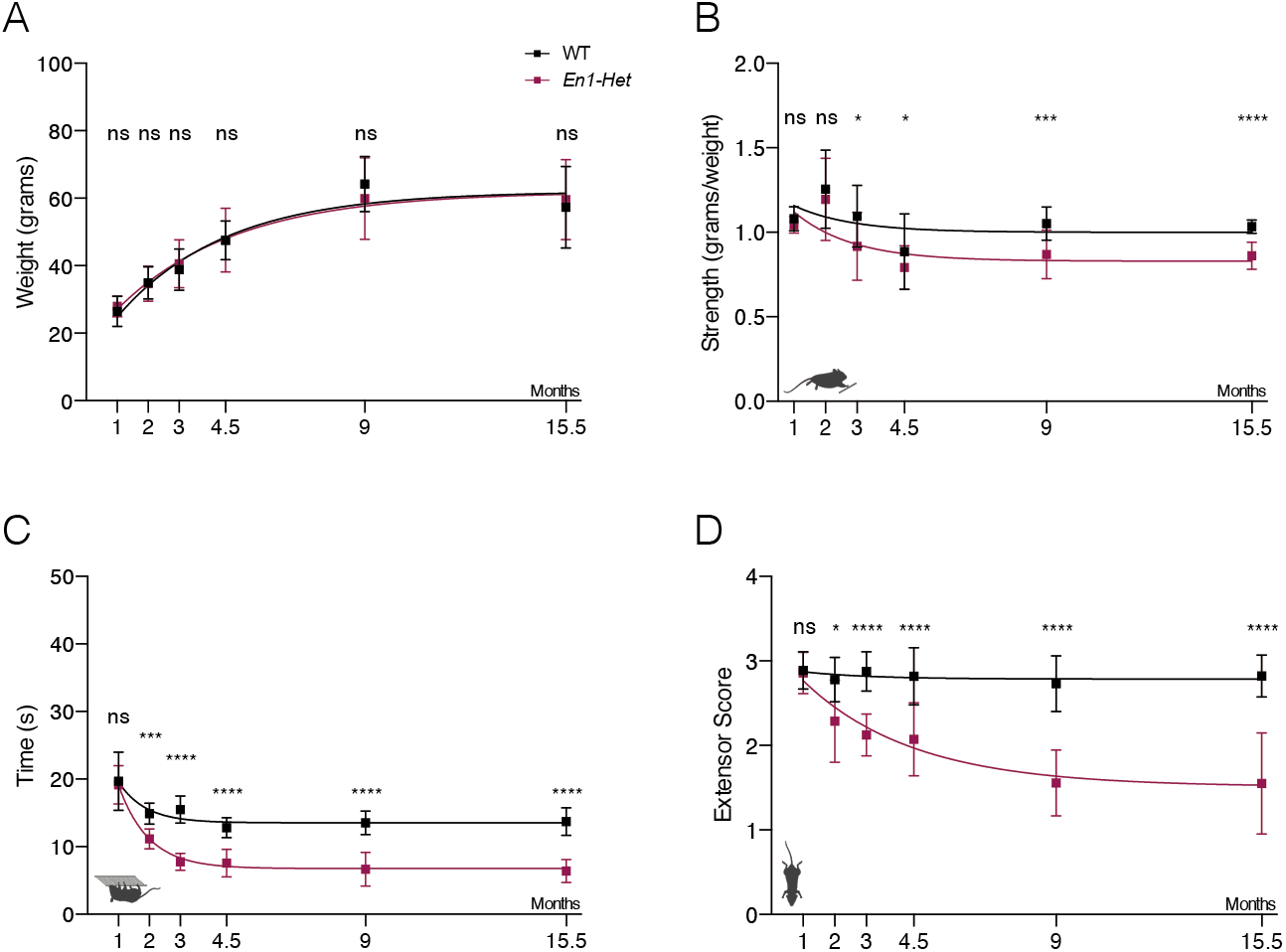
*En1-Het* mice develop muscle weaknessabnormal spinal reflex. **A**. Body weights of *En1-Het* and WT littermates are identical. **B**. *En1-Het* mice show reduced normalized grip strength starting at 3 months of age. **C**. *En1-Het* mice hold less time to the inverted grid, starting at 2 months of age. **D**. *En1-Het* mice show an abnormal hind limb reflex score, starting at 2 months of age. Unpaired T-test with equal SD comparing WT with *En1-Het* at each time point (*p<0.05; **p<0.005; ***p<0.0005; ****p<0.0001). n= 5 to 20.

We analyzed NMJs in three muscle groups with different vulnerabilities in motoneuron diseases (*15*). In the lumbrical muscle fibers no changes were found in the average number of AChR clusters between WT and *En1-Het* mice (Fig. 3A) but motor endplate area was reduced in *En1-Het* mice at 15.5 months (Fig. 3B). The percentage of endplates with perforations is normal in *En1-Het* mice through 9 months of age but significantly reduced at 15.5 months (Fig. 3C). Finally, the percentage of fully innervated endplates (neurofilament immunoreactivity overlaying more than 80% of the endplate) is significantly reduced in the *En1-Het* mice NMJ, starting at 3 months and worsening at 4.5 months (Fig. 3D). No differences between the two genotypes were observed in any of the NMJ morphological characteristics in extraocular and tongue muscles (Supplemental Fig. 3).

**Figure 3.**
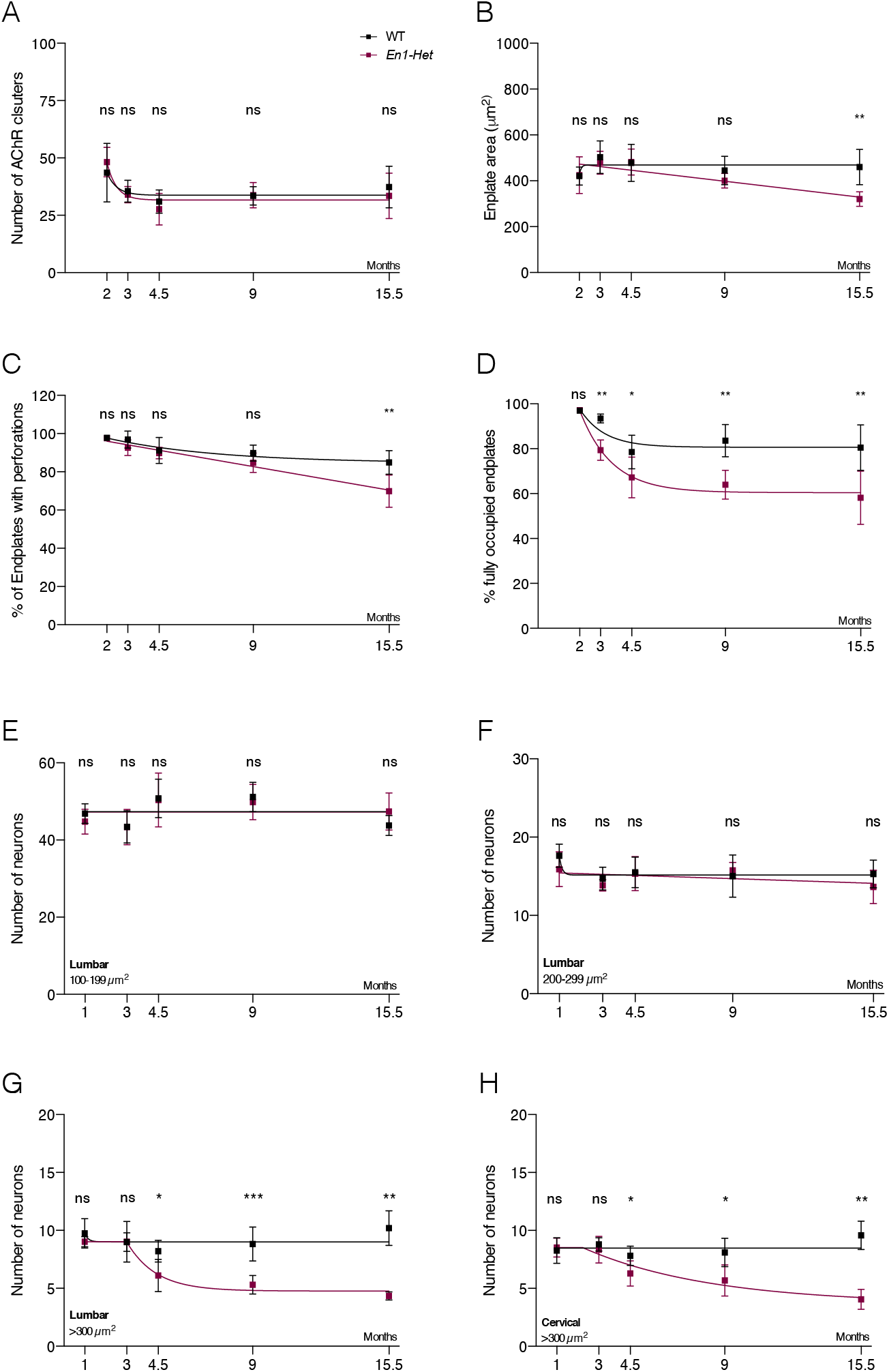
NMJ morphology and spinal neuron alterations changes in *En1-Het* mice. **A**. Lumbrical muscles of *En1-Het* and WT mice show similar number of AChR clusters. **B**. Endplate area is reduced in the *En1-Het* mice at 15.5 months of age. **C**. At 15.5 months, but not earlier, *En1-Het* mice have a reduction in the percentage of endplates with perforations. **D**. *En1-Het* mice show a decrease in the percentage of fully occupied endplates starting at 3 months of age. **E-H**. The number of Cresyl-violet-stained cells between 100 and 199 μm^2^ (**E**) or 200 and 299 μm^2^ (**F**) are similar in *En1-Het* and WT mice. The number of large lumbar (**G**) and cervical (**H**) αMNs neurons > 300 μm^2^ decreases in *En1-Het* mice starting at 4.5 months with a nearly 50% loss at 15.5 months. Unpaired T-test with equal SD comparing WT with *En1-Het* at each time point (*p<0.05; **p<0.005; ***p<0.0005; ****p<0.0001). n= 4 to 8.

Cresyl-violet stained neurons in the cervical and lumbar enlargements were classified by surface area (100-199 μm^2^, 200-299 μm^2^, and > 300 μm^2^). There was no change in the number of small and medium size neurons as shown for lumbar enlargement in Figure 3E,F. The loss of large Cresyl violet-stained neurons in the *En1-Het* mouse in cervical and lumbar enlargements was first observed at 4.5 months (≈20%) and reached ≈50% loss at 15.5 months (Fig. 3G,H). Choline acetyl transferase (ChAT) is selectively expressed in γ and αMNs, distinguishable by their size (*16*). When ChAT immunoreactivity was analyzed at 9 months, *En1-Het* spinal cord showed a maintenance of 200-299 μm^2^ neurons and a loss of large > 300 μm^2^ ChAT+ cells, with identical results for Cresyl violet-stained neurons (Supplemental Fig. 1 B-D). Thus, *En1-Het* mice show progressive αMNs loss, starting at 4.5 months and reaching about 50% at 15.5 months and this loss is not paralleled by an increase in intermediate size ChAT-stained MNs, precluding αMN shrinkage.

### Recombinant hEN1 reverses the *En1-Het* phenotype

A single injection of exogenous HPs rescue adult neurons from an *in vivo* stress for several weeks (*10*, *17*). We thus tested the effect of a single intrathecal hEN1 injection into the spinal cord of *En1-Het* mice. Since muscle weakness and NMJ abnormalies appear between 2 and 3 months and αMN loss at only 4.5 months, 3 months was selected as a reasonable injection time. Injection of 1μg of hEN1 was followed 24h later by strong EN1 immunofluorescence in neurons of the lumbar and cervical enlargements where EN1 was present in many ChAT-positive αMNs (Fig. 4A). The number of hEN1-stained cells was lower in the cervical region, probably due to its distance from the L5 injection site.

**Figure 4.**
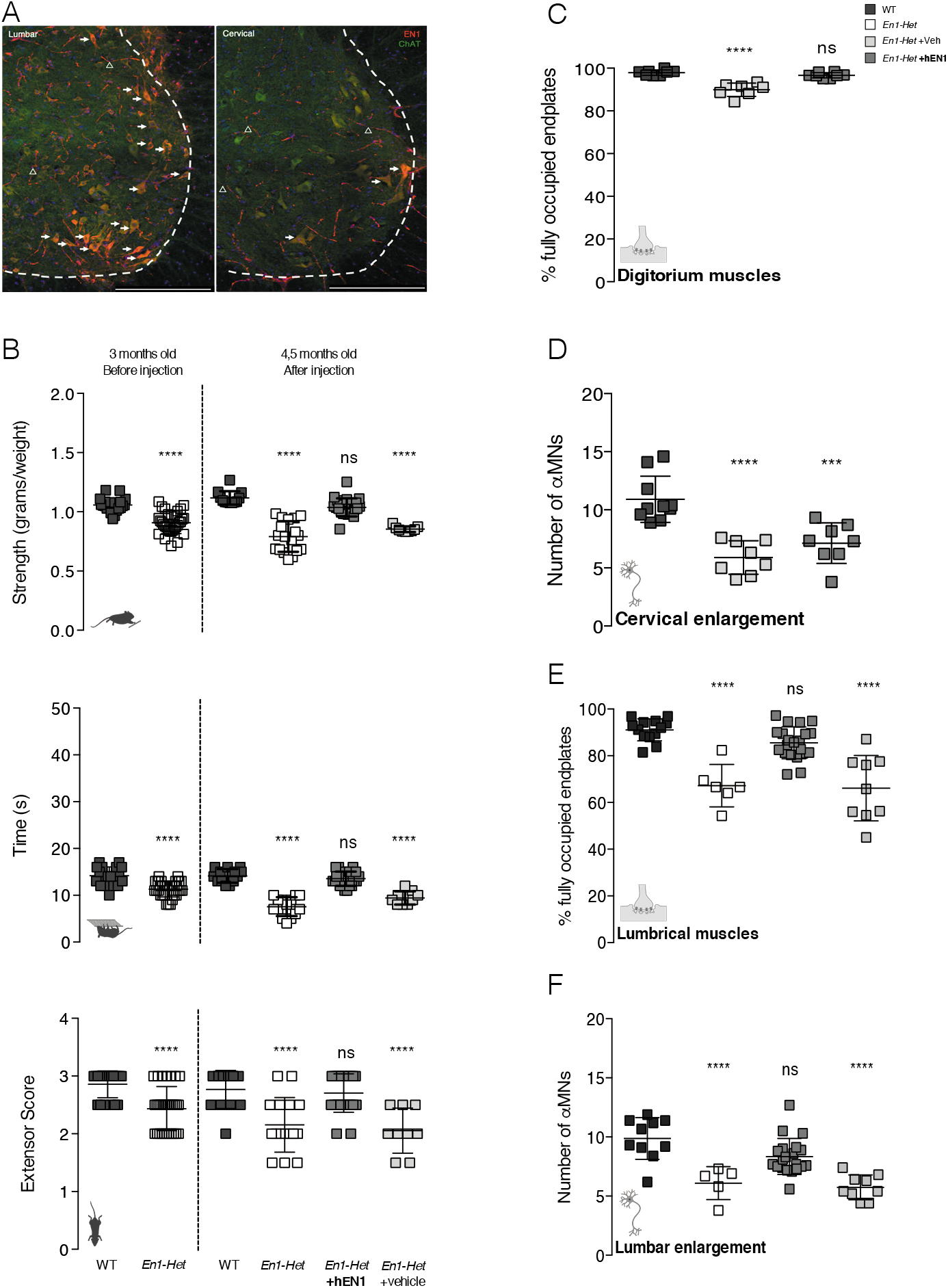
Intrathecal hEN1 restores *En1-Het* mice phenotype. **A**. Twenty-four hours after intrathecal injection of 1 μg of hEN1 at the L5 level, EN1 (red) and ChAT (green) are visualized in the lumbar (left) and cervical (right) enlargements. Arrows show internalized hEN1. Arrowheads show hEN1 in and/or around vessels. hEN1 injected at the lumbar level reaches ChAT-positive cells (arrows) in the cervical region (right). Scale bar: 100μm. **B**. Before hEN1 injection, 3-month-old *En1-Het* mice have reduced forepaw grip strength (top), reduced ability to hold onto the inverted grid (middle) and an abnormal hind limb extensor score (bottom). Injection of hEN1 at 3 months restores all scores to normal in 4.5-month-old *En1-Het* mice. **C, D** *En1-Het* mice injected at 3 months with vehicle and analyzed at 4.5 months show fewer fully innervated endplates (**C**) and fewer αMN (**D**); those injected with hEN1 have a percentage of fully occupied endplates similar to WT (**C**) with no αMN rescue in cervical spinal cord (**D**). **E, F.** *En1-Het* mice untreated or injected at 3 months with vehicle and analyzed at 4.5 months show a significant loss of fully occupied endplates and of αMNs, while those treated with hEN1 show normal endplate occupation (**E**) and a complete protection of lumbar αMNs (**F**). One-way ANOVA followed by a post hoc Dunnett’s test for comparisons to WT (*p<0.05; **p<0.005; ***p<0.0005; ****p<0.0001). n= 7 to 22.

Muscle strength and hindlimb extensor reflex were evaluated at 3 months of age and, 2 days later, *En1-Het* mice were separated into untreated, vehicle-injected or hEN1-injected groups. As above, 3-month-old *En1-Het* mice had reduced grip strength, reduced time holding onto the inverted grid and a lower hindlimb reflex score before intrathecal injections (Fig. 4B). One and a half months later, untreated *En1-Het* mice and those treated with vehicle continued to deteriorate in the three tests whereas the performance of hEN1-injected mice was indistinguishable from those of WT controls (Fig. 4B).

In 4.5-month-old *En1-Het* mice injected at 3 months with hEN1, the percentage of fully occupied endplates returns to normal at hindlimb and forelimb levels (Fig. 4C,E). Three-month-old *En1-Het* mice have a normal number of αMNs, but some loss is observed 1.5 months later. Figure 4F illustrates that hEN1 injected at 3 months prevents lumbar, but not cervical, αMN death. As a control, hEN1Q50A in which glutamine in position 50 of the homeodomain was mutated into alanine and can be internalized but lacks high-affinity binding to DNA (*18*) does not rescue muscular strength, extensor reflex, endplate occupancy or αMN numbers (Supplemental Fig. 4).

These experiments demonstrate that a single hEN1 lumbar injection at month 3 restores muscle strength, normalizes NMJ morphology and blocks lumbar, but not cervical αMN degeneration, for at least 1.5 months. Interestingly, while *En1-Het* mouse strength is restored in the forepaw grip strength and inverted grid tests, the survival of cervical αMNs is not rescued by hEN1. Because these tests involve the anterior limbs, it is likely that hEN1 had positive effects on the physiological activity of surviving αMNs.

To investigate if rescue could last more than 1.5 months, 3-month-old *En1-Het* mice injected with hEN1 were assessed weekly during the first month post-injection and monthly up to six months. Before injection, 3-month-old *En1-Het* mice showed significant muscle weakness and abnormal reflex compared to WT mice (Fig5. A-C, left panels). One week after injection, grip strength and extensor reflex had returned to WT values that were fully maintained for 3 months, and then performance declined to pre-injection levels. The improvement in the inverted grid test was more modest but followed a similar time-course. At 9 months, *En1-Het* mice injected with hEN1 at 3 months performed better in the 3 tests than age-matched nontreated *En1-Het* mice. *En1-Het* mice treated with hEN1 also showed higher NMJ innervation at 9 months than untreated age-matched *En1-Het* mice, although less than WT littermates (Fig. 5D, left panel). Partial rescue at 9 months was also observed for lumbar αMN survival (Fig.5D, right panel).

**Figure 5.**
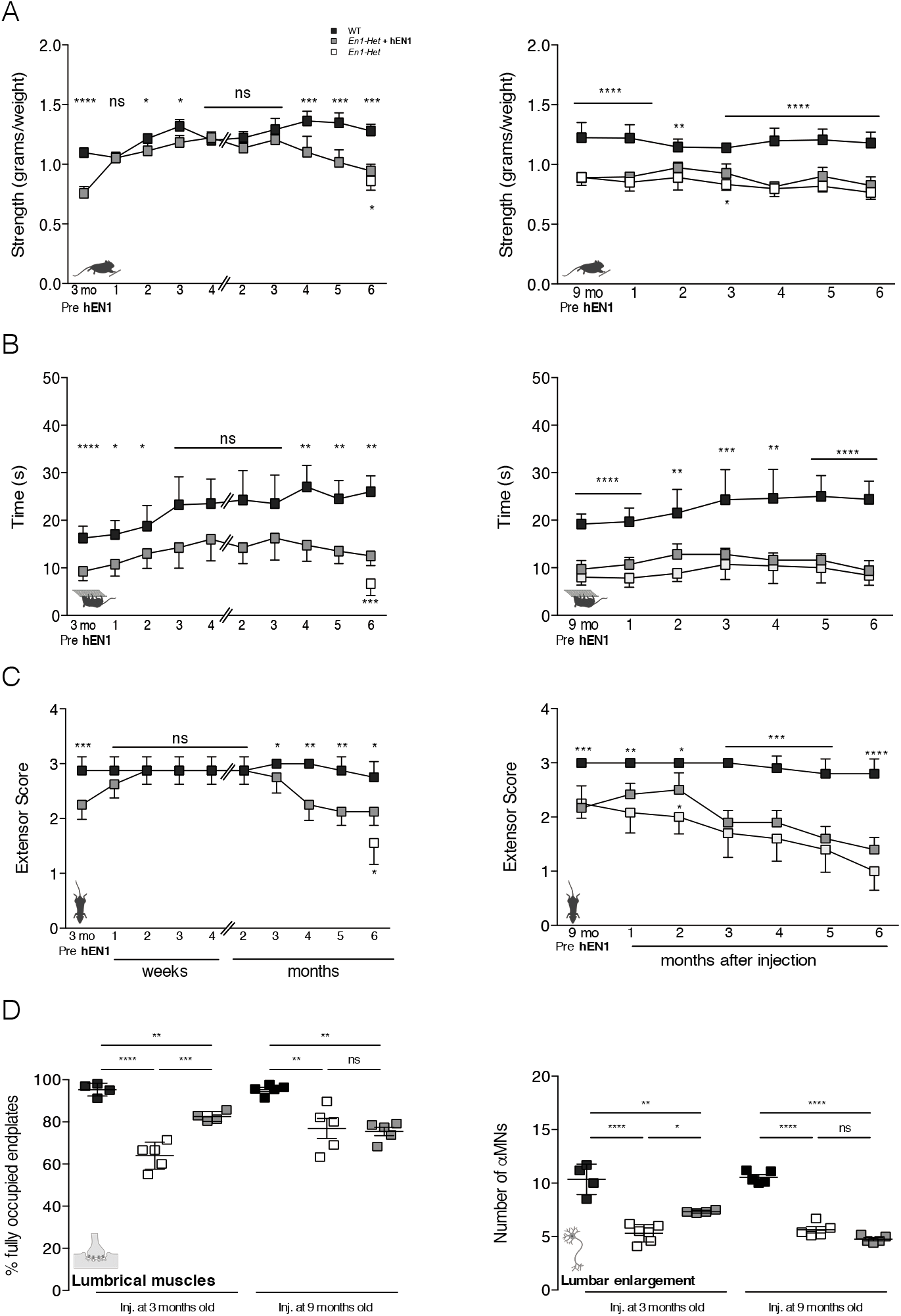
Intrathecal hEN1 has long-lasting rescuing activity. **A**. *Left*: Reduced grip strength observed in *En1-Het* mice returns to control levels one week after hEN1 injection at 3 months. This improvement lasts for 3 months before grip strength weakens. Grip strength remains higher in injected than in non-injected mice (open square) even at 9 months. *Right*: *En1-Het* mice treated at 9 months show no recovery. **B**. *Left*: Recovery in the inverted grid test starts at 3 weeks (no significant difference with WT siblings), lasts for 3 months and then declines. Strength remains higher in injected than in noninjected mice (open square) even at 9 months. *Right*: *En1-Het* mice treated at 9 months show no recovery. **C**. *Left*: Extensor reflex recovery starts at 2 weeks, lasts for 2 months and declines. The score remains higher in injected than in non-injected mice (open square) even at 9 months. *Right*: *En1-Het* mice treated at 9 months show no recovery. **D** *Left*: hEN1 injection at 3 months, but not at 9 months, partially restores endplate innervation pattern (fully occupied endplates) analyzed 6 months later while hEN1 injection at 9 months does not. *Right*: hEN1 injection at 3 months, but not at 9 months, partially maintains the number of lumbar αMNs analyzed at 9 months. Unpaired T-test with equal SD (*p<0.05; **p<0.005; ***p<0.0005; ****p<0.0001). n=4 to 9.

Finally, EN1 or vehicle were injected in 9-month-old *En1-Het* mice that manifested significant reductions in forepaw grip strength, time holding onto the inverted grid and extensor score (Fig. 5A-C, right panel). Compared to WT siblings, vehicle and hEN1-injected *En1-Het* mice remained significantly impaired in the three tests, with only a very limited improvement of the hEN1-treated mice, two months post injection, for the grip strength and extensor reflex tests. Anatomical analysis at the end of the experiment (15.5-month-old mice) showed no improvement in NMJ occupancy and αMN survival (Fig. 5D).

### hEN1 regulates the autophagy-associated p62 protein

EN1 protects mDA neurons from the SNpc. To investigate if similar protective pathways are activated by EN1 in mDA and motor neurons, we re-analyzed the transcriptome of SNpc lysates from adult WT and *En1-Het* mice (*10*). Differentially expressed genes (threshold p<0.05) were compared with a transcriptome analysis of WT MNs (*19*), and STRING analysis produced 402 candidate genes. Introducing in the gene interaction analysis 4 genes mutated in the most frequent forms of ALS: *Sod1*, *Fus*, *Tardbp-43* and C9orf72 yielded 20 genes of interest in the context of αMN physiology and survival.

*SQSTM1*/*p62* an autophagy-related gene with variants in 486 patients with familial ALS (*20*) was found to interact with the 4 ALS genes. To examine if expression of the gene is regulated by EN1 within αMNs, p62 levels were measured in large ChAT expressing neurons of WT and *En1-Het* mice spinal cords injected with hEN1Q50A or hEN1. Injections were made at 3 months and analyzed at 4.5 months. Compared to WT mice, *En1-Het* mice injected with the inactive form of EN1 show enhanced expression of p62 while the active transcription factor reduced this expression to WT levels, suggesting that hEN1 regulates p62 activity (Figure 6).

**Figure 6.**
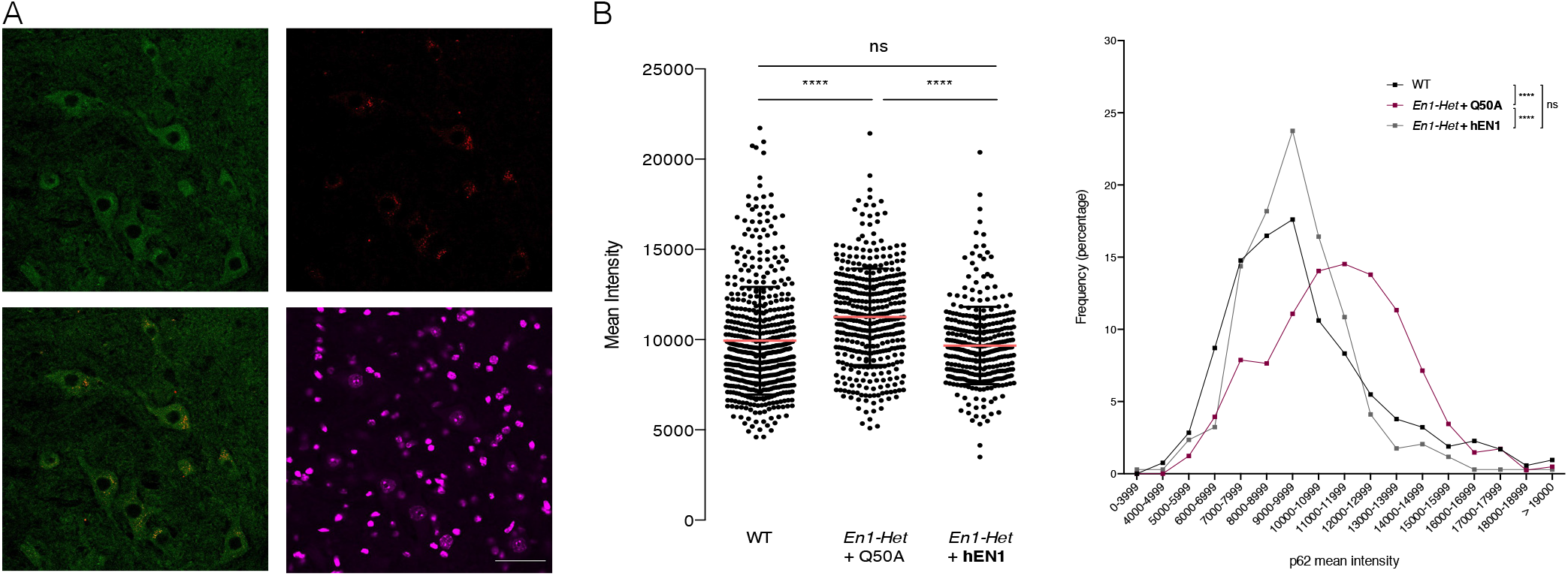
The autophagy marker p62 is regulated by EN1 in αMNs. **A.** Double-immunostaining for SQTSM1/p62 (red) and ChAT (green) shows SQTSM1/p62 expression in αMNs in 4.5-month-old *En1-Het* mice. Scale bar: 50 μm. **B.** *Left*. Quantification of SQTSM1/p62 intensity in large ChAT-positive αMNs demonstrates that SQTSM1/p62 expression is increased at 4.5 months in *En1-Het* mice injected at 3 months with hEN1Q50A and returns to normal in 4.5-month-old *En1-Het* mice injected with hEN1. One-way ANOVA (****p<0.0001) n=341 to 528 **B.** *Right*. The distribution of SQTSM1/p62 staining intensity in large αMNs shows that the shift of peak of intensity in *En1-Het* mice injected with hEN1Q50A is normalized in *En1-Het* mice injected with hEN1 Kolmogorov–Smirnov test (*p<0.05; ****p<0.0001)

## DISCUSSION

This study establishes the importance of non-cell-autonomous EN1 for αMN physiology and survival *in vivo*. In the *En1-Het* mouse, a single intrathecal injection has rapid and long lasting trophic activities. One day after injection, hEN1 accumulation in neurons, in particular αMNs, seems predominant, not precluding internalization by other neuronal or non-neuronal cell types.

Preferential hEN1 capture by αMNs suggests that they express EN1-binding sites, in line with the non-cell autonomous functions of EN1/2 reported earlier. This is reminiscent of OTX2, another HP that specifically accumulates in parvalbumin interneurons of the cerebral cortex (*3*) following its specific binding to glycosaminoglycans (GAGs) via a GAG binding domain (RKQRRERTTFTRAQL) (*21*). Similar GAG-binding domains are present in many HPs (*1*, *2*), including EN1 and EN2 for which this putative GAG-binding sequence is KEDKRPRTAFTAEQL. The potential role of this domain in the specific targeting of EN1 and the presence of EN1-binding sites at the αMN surface will be investigated in future studies.

In the *En1-Het* mouse, decreased strength between 2.5 and 3 months corresponds to a decrease in fully occupied endplates first observed at 3 months and precedes αMN loss first observed at 4.5 months. This suggests retrograde degeneration, with muscle strength and endplates affected before αMN loss. Similarly, in this *En1-Het* mouse *En1*-expressing mDA neuron terminals in the striatum show signs of degeneration at 4 weeks, while the neurons start dying at 6 weeks (*22*). In this context, the finding that hEN1 injection at 3 months not only prevents lumbar αMN death for months, but also restores normal endplate innervation and muscle strength indicates that neurodegeneration is slowed or halted for a considerable period of time.

αMN degeneration is rapid between 3 and 4.5 months and then slows until at least 15.5 months when ≈40% αMNs remain. This heterogenity of αMNs for their dependency on EN1 is reminiscent of the lesser resistance of mDA neurons in the SNpc than in the Ventral Tegmental Area to *En1* hypomorphism (*7*). It is also of note that the amounts of hEN1 reaching the cervical enlargement, while not sufficient to prevent αMN death, restored forelimb muscle strength and endplate innervation, suggesting that EN1 has beneficial effects on the remaining αMNs. Whether this involves sprouting of αMN terminals to occupy empty innervation sites will be analyzed.

Homeoproteins act cell- and non-cell-autonomously. A strictly cell autonomous activity in V1 interneurons would mean that they are dysfunctional in the *En1-Het* mouse. Deficits in inhibitory interneurons in the spinal cord of MN disease models have been reported and Renshaw cell pathology has been observed following αMN loss (*23*, *24*). Not mutually exclusive, EN1 might act in a non-cell autonomous way. Firstly, it may regulate other trophic factors to support αMN survival. Secondly, it might be secreted and internalized by intermediate cell types, such as astrocytes or microglial cells, enhancing their αMN-directed trophic activity (*25*, *26*). Finally, EN1 may transfer from V1 interneurons to αMNs to exert a direct protective activity as observed for OTX2 (*17*).

Neutralizing extracellular EN1 with an anti-EN1/2 scFv reduces endplate innervation and muscle strength without αMN loss, at least 4 months following viral injection. While longer survival times may show αMN loss, we cannot exclude that extracellular EN1 acts in conjunction with other factors with decreased secretion by *En1-Het* interneurons. Extracellular ENGRAILED co-signaling has been shown with DPP in the fly wing disk (*14*) and with EphrinA5 in the chick optic tectum (*5*). Mechanisms for protection of mDA neurons by a single EN1 injection have been characterized. Among them are the regulation of Complex I mitochondrial activity, DNA break repair, heterochromatin maintenance, and the repression of long interspersed nuclear element (LINE-1) expression (*8*, *10*, *11*, *27*). The activation of these mechanisms will be the object of future studies.

The *En1-Het* mouse presents phenotypes highly reminiscent of changes in ALS patients and in many ALS mouse models. However, although they may exist in some sporadic cases, no ALS phenotype has been associated with mutations affecting *EN1* expression, precluding that the *En1-Het* mouse be considered as an ALS model. Yet, *En1* might interact genetically with pathways leading to ALS, and EN1 could be envisaged as a long-lasting therapeutic protein, as shown in classical mouse and non-human primate models of Parkinson Disease (*28*).

Strikingly, the bioinformatic analysis led to a list of only 20 genes that interact with genes mutated in fALS. One of them, *SQSTM1/p62*, a known regulator of autophagy (*28*), associates with the four fALS genes included in our analysis and is up-regulated in 4.5-month-old *En1-Het* mice injected at 3 months with an inactive form of EN1, but is normal in *En1-Het* mice injected with active EN1. The ensuing hypothesis that *En1* may interact with fALS genes will be evaluated by generating double heterozygotes for *En1* and fALS genes.

## SUPPLEMENTAL INFORMATION

### MATERIALS AND METHODS

#### Animal Management

All animal treatments followed the guidelines for the care and use of laboratory animals (US National Institutes of Health), the European Directive number 86/609 (EEC Council for Animal Protection in Experimental Research and Other Scientific Utilization) and French authorizations n° 00703.01 “Therapeutic homeoproteins in Parkinson Disease” and n° APAFIS #6034-2016071110167703 v2, “Spinal cord motoneuron neuroprotection” delivered by the Minister of higher education, research and innovation. Adult mice were housed two to five per cage with *ad libitum* access to food and water and under a 12h light/dark cycle. Transgenic mouse strain *En1-Het* was bred by the Rodent Breeding Services provided by the Animal Care Services at College de France. Females and males were included in all studies. The endpoint limit for euthanasia was a 15% or greater loss of bodyweight or signs of paralysis; in all experiments, no mouse reached these endpoints.

#### Behavioral analyses

Mice were habituated to the behavioral room and to the experimenter 24 hours before the day of testing and again before each behavioral test. All tests were performed on the same day and behavioral assessment was carried by evaluators blind to genotype and treatment.

##### Forepaw Grip Strength

The Transducer (IITC Life Science Grip Strength Meter, ALMEMO 2450 AHLBORN, World Precision Instruments) was calibrated and the scale of values set to grams. Each mouse was lifted by the tail to position the front paws at the height of the bar (about 15 cm) and then moved towards the bar. When symmetric firm grip with both paws was established the mouse was pulled backward at a constant speed until the grasp was broken and the maximal value was recorded. The test was repeated 5 times per animal with a minimal resting time of 5 minutes between tests and the mean of all values was normalized to the weight of each animal.

##### Inverted Grid Test

The mouse was placed on a wire grid (15×10 cm) and allowed to explore it. After 3-5 minutes, the grid was raised 30 cm above a soft surface and slowly inverted. Latency to release was recorded three times per mouse with a minimum resting time of 5 minutes between trials. The longest latency was used for analysis.

##### Hindlimb Extensor Reflex

Mice were suspended by the tail at a constant height (30 cm) and scored for hindlimb extension reflex. The scores were assigned from 0 to 3 as follows: 3 for normal symmetric extension in both hind limbs without visible tremors; 2.5 for normal extension in both hind limbs with tremor in one or both paws; 2.0 for unequal extension of the hind limbs without visible tremors; 1.5 for unequal extension in the hind limbs with tremors in one or both paws, 1.0 for extension reflex in only one hindlimb, 0.5 for minimum extension of both hindlimbs, 0 for absence of any hindlimb extension.

#### Tissue preparation

##### Spinal cord

Adult mice were euthanized by a 1 μl/g body weight dose of Dolethal (Euthasol: 4 μg/μl). Spinal cords were dissected and placed in Phosphate Buffer Saline (PBS) to remove meninges and surrounding connective tissues. Cervical and lumbar enlargements were separately placed in paraformaldehyde 4% (PFA, Thermo Scientific) for 1 h at room temperature (RT) with gentle mixing, washed in PBS three times for 30 minutes at RT and placed in PBS with 20% sucrose overnight at 4°C. After cryoprotection, the tissue was embedded in Tissue Freezing Medium (TFM, Microm Microtech), frozen on dry ice and 30 μM sections prepared using an HM 560 Microm cryostat (Thermo Scientific).

##### Muscle

The extraocular muscle (EOM), tongue and lumbrical muscles were dissected in cold PBS and fixed at RT in 4% PFA for 10 minutes for extraocular and lumbrical muscles (1 hour for the tongue) washed in PBS and cryoprotected. Extraocular and lumbrical muscle wholemounts were processed *in toto* to visualize the entire innervation pattern and detailed analysis of the neuromuscular junctions (*16*). Tongue muscles were embedded and sectioned at 30 μm.

##### Cresyl violet staining

Slides with 30 μm spinal cord sections were washed in PBS 3 times, cleared in O-Xylene (CARLO-HERBA) for 5 minutes, then hydrated in graded alcohols with increasing water and placed in Cresyl Violet acetate (MERCK). Sections where then dehydrated in increasing alcohol and mounted in Eukitt^®^ Quick-hardening mounting medium (Sigma).

##### Spinal cord and muscle immunofluorescence labeling

Slides with 30 μm spinal cord or muscle sections were washed in PBS and permeabilized with 2% Triton. After 30 minutes at RT in 100 μM glycine buffer, sections were blocked in 10% Normal Goat Serum (NGS, Invitrogen) or Fetal Bovine Serum (FBS, Gibco) in the presence of 1% Triton and incubated with primary antibodies (sheep anti-Choline Acetyltransferase (ChAT) ABCAM 1:1000, goat anti-ChAT Millipore 1:500, rabbit anti-EN1 1:300 (*7*), mouse anti-neurofilament 165kDa DSHB 1:50, mouse anti-synaptic vesicle glycoprotein 2A DSHB 1:100 and rabbit anti-p62 ABCAM 1:1000) overnight at 4°C, washed and further incubated with secondary antibodies for 2 hours at RT. For muscle staining, α-bungarotoxin (Alexa Fluor 488 conjugate) was included at the same time as the secondary antibodies. Slides were washed and mounted with DAPI Fluoromount-G^®^ (Southern Biotech). Controls without primary antibodies were systematically included.

#### RT-qPCR

Spinal cords were removed as above and lumbar enlargements were rapidly frozen on dry ice. Total RNA was extracted (RNeasy Mini kit, Qiagen) and reverse transcribed using the QuantiTect Reverse Transcription kit (Qiagen). RT-qPCR was done using SYBR-Green (Roche Applied Science) and a Light Cycler 480 (Roche Applied Science). Data were analyzed using the “2-ddCt” method and values were normalized to *Glyceraldehyde 3-phosphate dehydrogenase* (*Gapdh*). The following primers were used: *Engrailed-1* sense: CCTGGGTCTACTGCACACG, antisense: CGCTTGTTTTGGAACCAGAT; *Gapdh* sense: TGACGTGCCGCCTGGAGAAAC, antisense: CCGGCATCGAAGGTGGAAGAG.

#### Protein production

Human EN1 (hEN1) was produced as described (*17*) and endotoxins were removed by Triton X-144 phase separation. In brief, pre-condensed 1% Triton X-144 (Sigma) was added to the protein preparation. The solution was incubated for 30 minutes at 4°C with constant stirring, transferred to 37°C for 10 minutes and centrifuged at 4000 rpm for 10 minutes at 25°C. The endotoxin-free protein was aliquoted and kept at −80°C.

#### Intrathecal injections

Mice were anesthetized with 1 μl/g ketamine-xylazine (Imalgene1000: 10 μg/μl, Ruompur 2%: 0,8 μg/μl) and placed on the injection platform. The tail of the animal was taken between two fingers and the spinal column was gently flattened against a padded surface with the other hand. The L3 vertebral spine was identified by palpation and a 23G × 1” needle (0.6×25 mm Terumo tip) was placed at the L1 and T13 groove and inserted through the skin at an angle of 20° (*16*). The needle was slowly advanced into the intervertebral space until it reached the injection point, provoking a strong tail-flick reflex. Five μl were injected at a rate of 1 μl/min with or without 1μg recombinant protein. The needle was left in place for two minutes after injection and then slowly removed. Animals were placed in a warmed chamber until recovery from anesthesia. Extensor reflex and gait analysis were examined 2 and 24 hours after injection to verify the absence of spinal cord damage.

#### RNAscope Fluorescent in Situ Hybridization

Mice aged 4.5 months were euthanized with Dolethal and transcardially perfused with 0.9% NaCl. The lumbar region of the spinal cord was dissected, fixed overnight in 4% PFA at 4°C and cryoprotected in PBS 30% sucrose for 48 hours at 4°C. Spinal cord sections 40 μm-thick were prepared using a Leitz (1400) sliding microtome. Sections were washed in PBS, incubated with RNAscope hydrogen peroxide solution from Advanced Cell Diagnostics (ACD) for 10 min at RT, rinsed in Tris-buffered saline with Tween (50 mM Tris-Cl, pH 7.6; 150 mM NaCl, 0.1% Tween 20) at RT, collected on Super Frost plus microscope slides (Thermo Scientific), dried at RT for 1 hour, rapidly immersed in ultrapure water, dried again at RT for 1 hour, heated for 1 hour at 60°C and dried at RT overnight. The next day, sections were immersed in ultrapure water, rapidly dehydrated with 100% ethanol, incubated at 100°C for 15 minutes in RNAscope 1X Target Retrieval Reagent (ACD), washed in ultrapure water and dehydrated with 100% ethanol for 3 minutes at RT. Protease treatment was carried out using RNAscope Protease Plus solution (ACD) for 30 minutes at 40°C in a HybEZ oven (ACD). Sections were then washed in PBS before *in situ* hybridization using the RNAscope Multiplex Fluorescent V2 Assay (ACD). Probes were hybridized for 2 hours at 40°C in HybEZ oven (ACD), followed by incubation with signal amplification reagents according to the manufacturer’s instructions. Probes were purchased from ACD: Mm-*En1*-C1 (catalog #442651), Mm-*Chat*-C2 (catalog #408731-C2), Mm-*Calb1*-C3 (catalog #428431-C3). The hybridized probe signal was visualized and captured on an upright Leica CFS SP5 confocal microscope with a 40x oil objective.

#### Bioinformatic analysis

Differentially expressed genes (cutoff p<0.05) from the RNA-seq performed on microdissected six-week-old SNpc from wild type (WT) and *En1*-*Het* mice (GSE72321 (*10*)) were compared to genes identified from an RNAseq of microdissected MNs from WT mice (GSE38820 (*19*)). Transcripts present in the two lists were retained. Using STRING database, 10 biological processes of interest were selected including regulation of gene expression, cellular response to stress, cell communication, regulation of I-kappaB kinase/NF-kappaB signaling, cellular response to DNA damage stimulus, response to oxygen-containing compounds, locomotion, DNA repair, modulation of chemical synaptic transmission, and brain development. Using these pathways reduced the list of selected genes to 402. The interaction of the selected genes with the four main genes involved in familial ALS (fALS): *Sod1*, *Fus, C9Orf72* and *Tardbp* was investigated.

#### Image analyses

Cresyl violet stained spinal cord section images were acquired with a Nikon-i90 microscope under bright field conditions at 10x with a Z-stack step of 0.5 μm. Immunofluorescence stained spinal cord sections images were acquired with a Leica SP5 inverted confocal microscope at 20x (Leica DMI6000) and acquisitions of 3D z-stack (0.5 μm) were made using the UV (405 nm, 50 mW), Argon (488 nm, 200 mW) and DPSS (561 nm, 15 mW) lasers. For cell quantification, at least five spinal cord sections separated by ≥ 900 μm were analyzed for each animal. ChAT+ cells and Cresyl violet stained cells with a soma area above 100 μm^2^ were manually outlined using ImageJ software (NIH, Bethesda, MD) and their area determined. Analyses were carried out on Z-stacks through the entire 30 μm thickness of the section. Cells were classified as small (100-199 μm^2^ cross sectional area), intermediate (200-299 μm^2^) or large (> 300 μm^2^). Supplemental Figure 1 illustrates that ChAT and Cresyl violet gave similar results for medium size (200-299 μm^2^) and large (>300 μm^2^) neurons thus allowing us to characterize αMNs on the basis of size (>300 μm^2^), with or without additional ChAT staining. For p62 staining analysis, the motoneuron region of five lumbar sections were acquired at 40x for each animal. The mean intensity was measured in ChAT+ cells with a soma greater than 300 μm^2^ using ImageJ software. For each experiment, image acquisition was performed with the same parameter settings of the confocal microscope to allow for comparison between experimental conditions.

Lumbrical, EOM and tongue muscles were imaged with a Leica SP5 inverted confocal microscope (Leica DMI6000) with a motorized XY stage. 3D z-stack (0.5 μm) acquisitions were made as above (UV, Argon and DPSS) and images analyzed using ImageJ software. Analyses were performed blind to the genotype and treatment. Endplates where categorized as fully innervated (neurofilament overlying more than 80% of the endplate), partially innervated (neurofilament overlying up to 80% of the endplate) or denervated (no neurofilament overlying the endplate. Endplate morphology was evaluated by counting the number of endplates with perforations (areas lacking α-bungarotoxin staining). For postsynaptic analysis, each endplate was manually outlined using ImageJ software and its area calculated. All analyses were performed on the entire Z-stacks through the NMJ.

#### Statistical analyses

Results are expressed as mean ± SD. Statistical significance was determined as follows. For RT-qPCR, WT and *En1-Het* mice were compared by Unpaired T-test with equal SD. For behavioral and NMJ analyses and αMN counting in the time-course study, groups were compared by Unpaired T-test with equal SD comparing WT with *En1-Het* for each time point. For intrathecal injections, behavioral and NMJ analyses and αMN counting, experimental data were compared by one-way ANOVA followed by a post hoc Dunnett’s test for comparisons to WT. For behavioral analysis in the time-course protection of injected hEN1, groups were compared by Unpaired T-test with equal variances comparing WT with *En1-Het* injected at each time point.

---

**Supplemental Figure 1.**
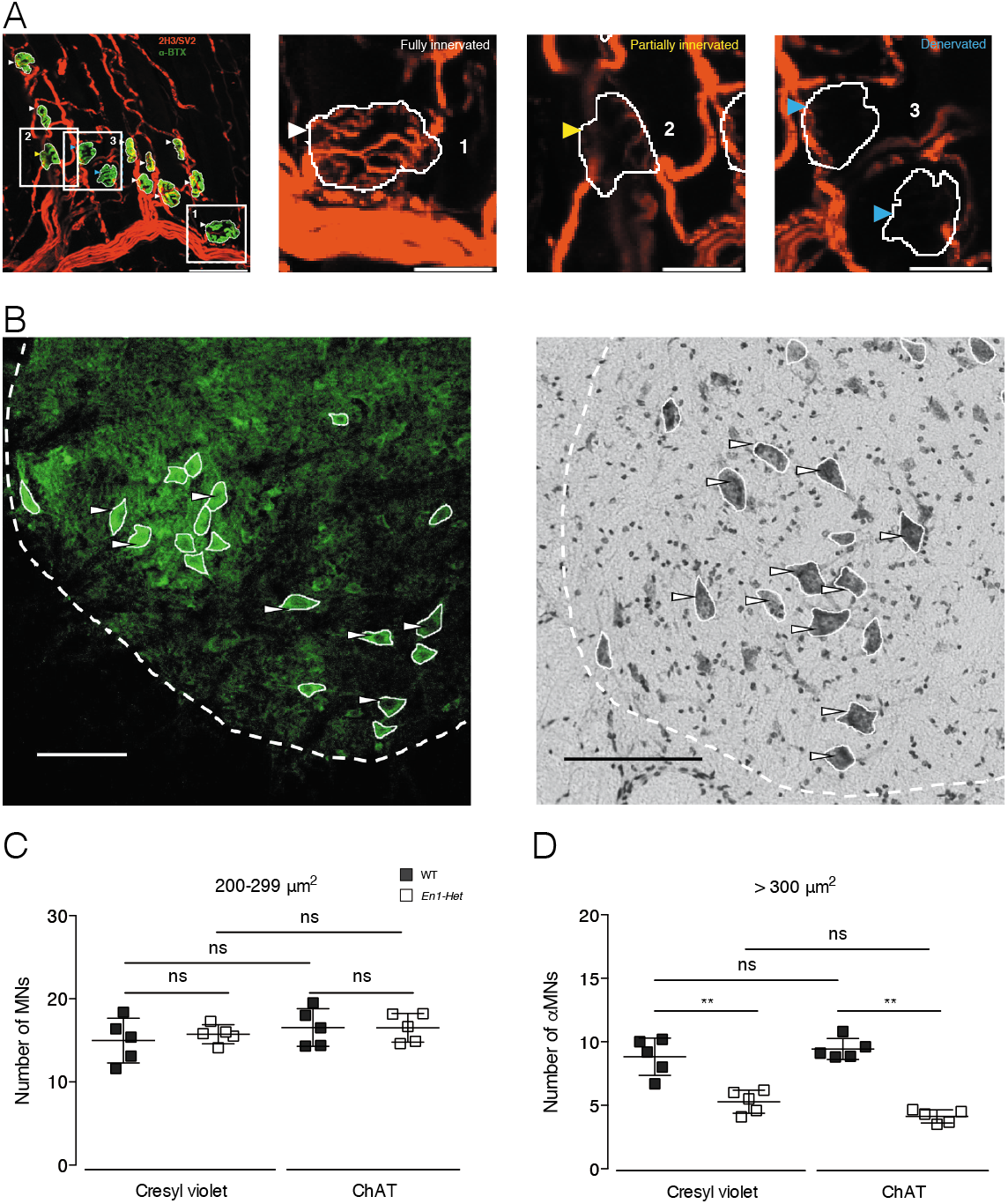
Identification of αMNs by ChAT staining. **A.** Representative confocal images of lumbrical muscle NMJ stained for neurofilament, synaptic vesicle glycoprotein 2A (2H3/SV2, red) and α-BTX (green). Endplate area (α-BTX+) was manually delimited (white profiles) and categorized as fully innervated (neurofilament overlying more than 80% of the endplate, white arrows), partially innervated (overlying up to 80% of the endplate, yellow arrow) or denervated (no neurofilament overlying the endplate, blue arrows). Scale bar: 50 μm left panel, 25 μm other panels. **B.** Representative confocal micrograph (left, ChAT staining) and light field micrograph (right, Cresyl violet) of the left ventral lumbar enlargement of 9-month-old WT mouse showing manually delimited areas (white profiles). Arrows show examples of cells with somas larger than 300 μm^2^. Scale bar: 100 μm. **C, D**. The number of ChAT+ cells between 200 and 299 μm^2^ (putative γMNs) is similar between genotypes (left) whereas that of cells with a surface > 300 μm^2^ (αMNs) is reduced in *En1-Het* mice. Cresyl violet and ChAT staining give similar results. Unpaired T-test with equal SD (*p<0.05; **p<0.005; ***p<0.0005; ****p<0.0001). n=5 for all groups.

**Supplemental Figure 2.**
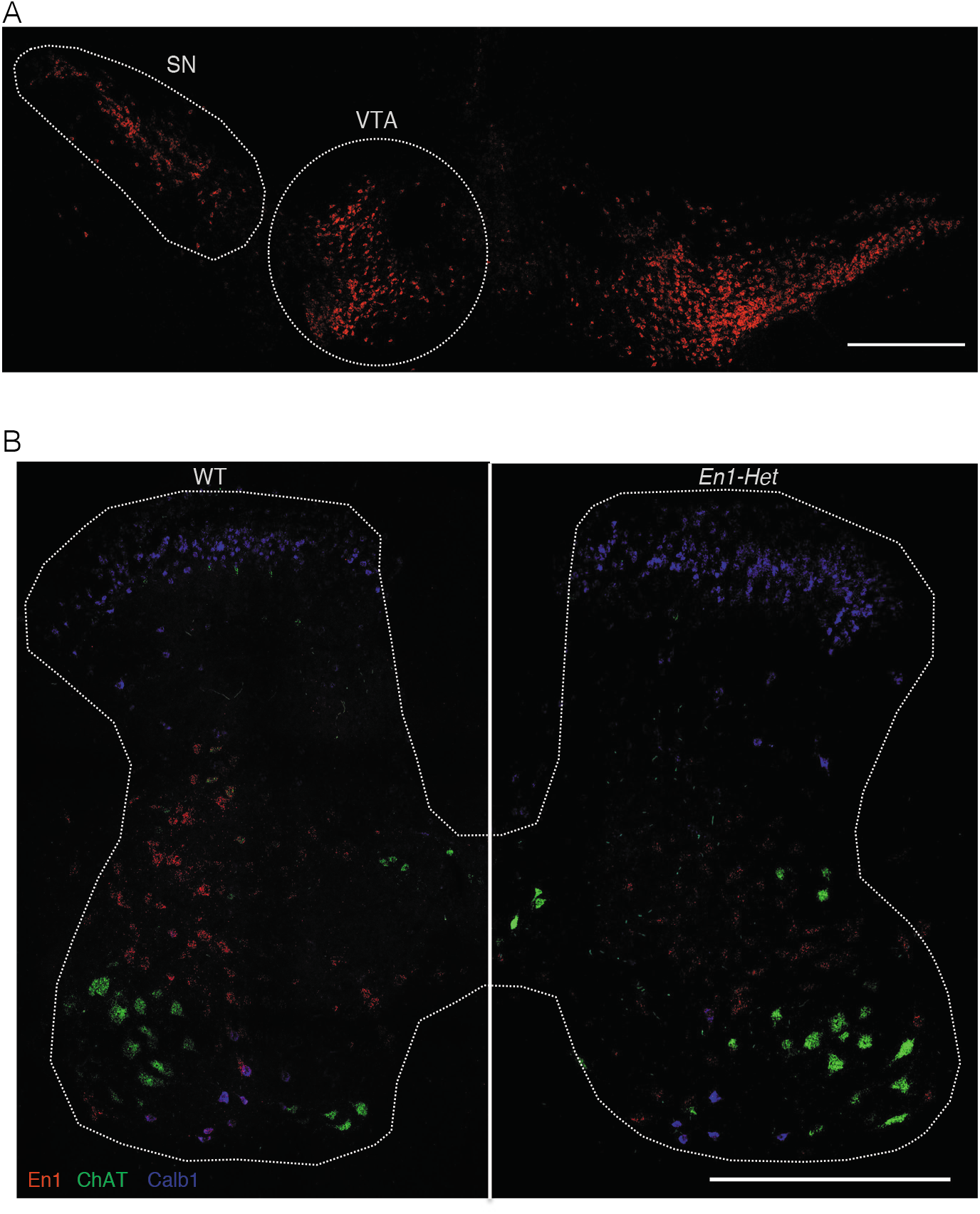
*Engrailed-1* expression in adult midbrain nuclei and spinal cord. **A.** RNAscope *in situ* hybridization in the adult brain shows strong and selective expression of *Engrailed-1* in the ventral tegmental area (VTA) and Substancia Nigra (SN) of the ventral midbrain. **B.** RNAscope triple *in situ* hybridization on lumbar spinal cord section shows the expression of *Engrailed-1* (red), *Choline acetyl transferase* (green) and *Calbindin-1* (blue) genes in WT (left) and *En1* heterozygote (right). Note the reduced *Engrailed-1* expression in the *En1* heterozygote. Scale bar: 500 μm

**Supplemental Figure 3.**
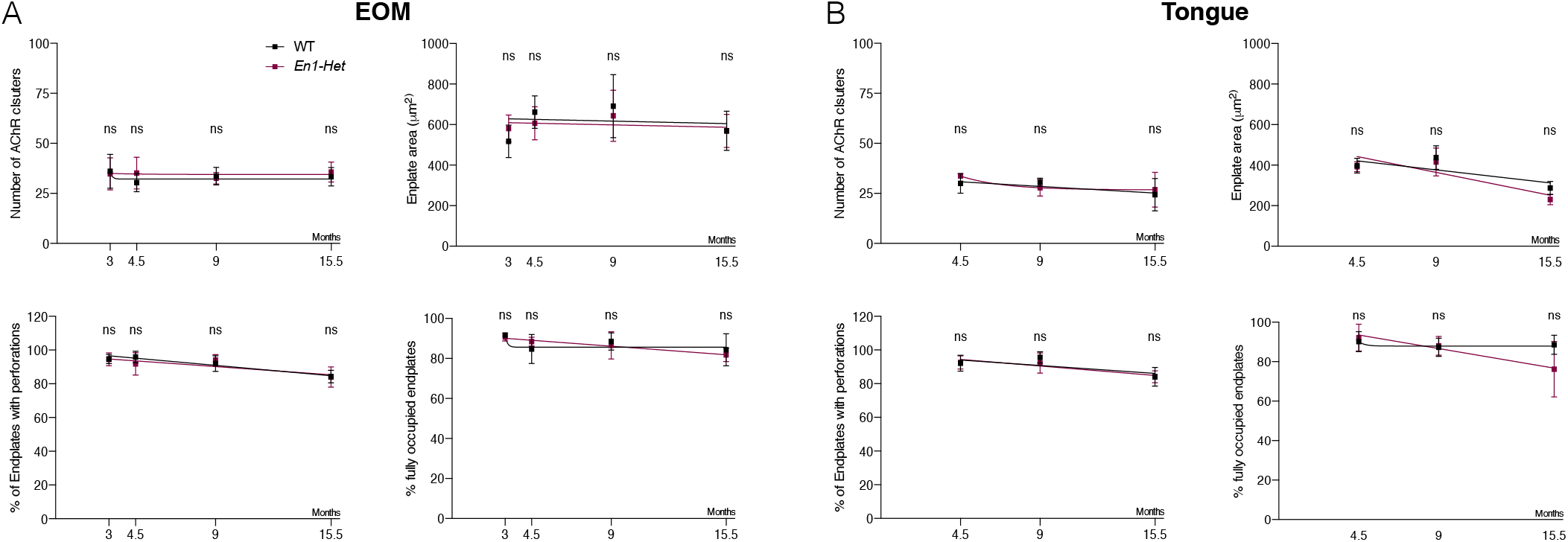
Analysis of EOM and tongue NMJs. **A**. In the EOM, the number of AChR clusters (top left), the size of endplate area (top right), the percentage of endplates with perforation (bottom left) and that of fully occupied endplates (bottom right) did not differ between *En1-Het* and WT mice. **B**. In the tongue, the number of AChR clusters (top left), the size of endplate area (top right), the percentage of endplates with perforation (bottom left) and that of fully occupied endplates (bottom right) did not differ between *En1-Het* and WT mice. Unpaired T-test with equal SD comparing WT with *En1-Het* at each time point (*p<0.05; **p<0.005; ***p<0.0005; ****p<0.0001). n= 4 to 8.

**Supplemental Figure 4.**
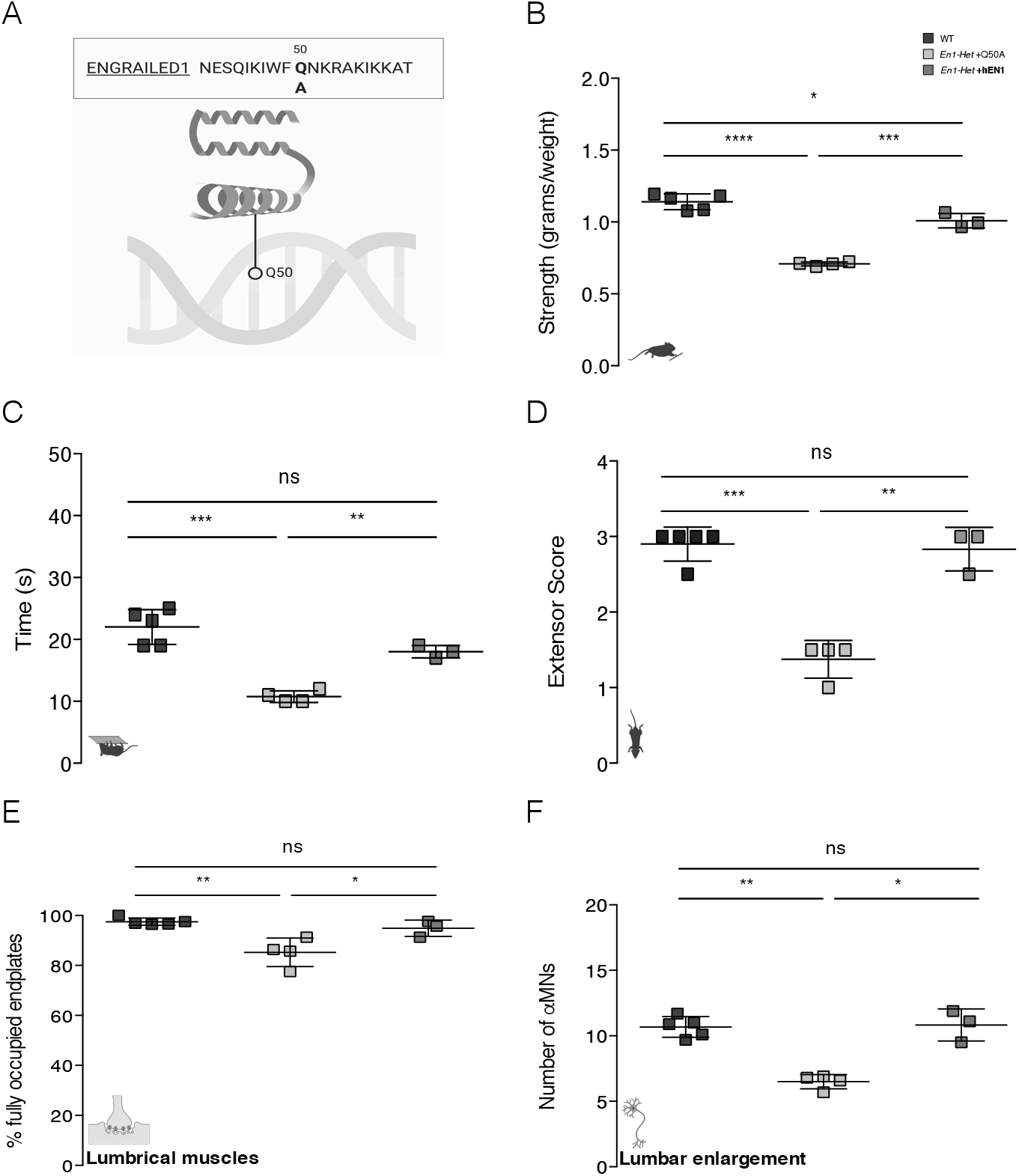
The hEN1Q50A mutant with reduced high affinity DNA binding does not reverse the *En1-Het* mouse phenotype. **A**. Scheme of the Q50A mutation in the EN1 homeodomain **B-F**. Compared to hEN1, hEN1Q50A has no rescuing activity in the anterior grip strength (**B**), time on the grid (**C**) and extensor reflex (**D**) tests. Accordingly, injection of the mutated protein does not protect against the decrease in endplate occupancy (**E**) or the loss of αMNs (**F**). Unpaired T-test with equal SD (*p<0.05; **p<0.005; ***p<0.0005; ****p<0.0001). n=3 to 5.

## Acknowledgements

We thank the Collège de France imaging/animal facilities, Dr. A.-C. Boulay for assistance in RNAscope, A.A. Di Nardo for careful reading of the manuscript and A. Bibimbou for technical assistance.

## References

1. A. Prochiantz, A. A. D. Nardo, Homeoprotein signaling in the developing and adult nervous system. Neuron. 85, 911–925 (2015).

2. A. A. D. Nardo, J. Fuchs, R. L. Joshi, K. L. Moya, A. Prochiantz, The Physiology of Homeoprotein Transduction. Physiol Rev. 98, 1943–1982 (2018).

3. S. Sugiyama, A. A. D. Nardo, S. Aizawa, I. Matsuo, M. Volovitch, A. Prochiantz, T. K. Hensch, Experience-dependent transfer of Otx2 homeoprotein into the visual cortex activates postnatal plasticity. Cell. 134, 508–520 (2008).

4. H. Kaddour, E. Coppola, A. A. D. Nardo, C. L. Poupon, P. Mailly, A. Wizenmann, M. Volovitch, A. Prochiantz, A. Pierani, Extracellular Pax6 Regulates Tangential Cajal–Retzius Cell Migration in the Developing Mouse Neocortex. Cereb Cortex (2019), doi:10.1093/cercor/bhz098.

5. A. Wizenmann, I. Brunet, J. S. Y. Lam, L. Sonnier, M. Beurdeley, K. Zarbalis, D. Weisenhorn-Vogt, C. Weinl, A. Dwivedy, A. Joliot, W. Wurst, C. Holt, A. Prochiantz, Extracellular Engrailed participates in the topographic guidance of retinal axons in vivo. Neuron. 64, 355–366 (2009).

6. N. Kim, K. W. Min, K. H. Kang, E. J. Lee, H.-T. Kim, K. Moon, J. Choi, D. Le, S.-H. Lee, J. W. Kim, Regulation of retinal axon growth by secreted Vax1 homeodomain protein. eLife. 3, e02671 (2014).

7. L. Sonnier, G. L. Pen, A. Hartmann, J.-C. Bizot, F. Trovero, M.-O. Krebs, A. Prochiantz, Progressive Loss of Dopaminergic Neurons in the Ventral Midbrain of Adult Mice Heterozygote for Engrailed1. J Neurosci. 27, 1063–1071 (2007).

8. D. Alvarez-Fischer, J. Fuchs, F. Castagner, O. Stettler, O. Massiani-Beaudoin, K. L. Moya, C. Bouillot, W. H. Oertel, A. Lombès, W. Faigle, R. L. Joshi, A. Hartmann, A. Prochiantz, Engrailed protects mouse midbrain dopaminergic neurons against mitochondrial complex I insults. Nature Neuroscience. 14, 1260–1266 (2011).

9. N. Thomasson, E. Pioli, C. Friedel, A. Monseur, J. Lavaur, K. L. Moya, E. Bezard, A. Bousseau, A. Prochiantz, Engrailed-1 induces long-lasting behavior benefit in an experimental Parkinson primate model. Movement Disorders. 34, 1082–1084 (2019).

10. H. Rekaik, F.-X. B. de Thé, J. Fuchs, O. Massiani-Beaudoin, A. Prochiantz, R. L. Joshi, Engrailed Homeoprotein Protects Mesencephalic Dopaminergic Neurons from Oxidative Stress. CellReports. 13, 242–250 (2015).

11. F. B. de Thé, H. Rekaik, E. Peze-Heidsieck, O. Massiani-Beaudoin, R. L. Joshi, J. Fuchs, A. Prochiantz, Engrailed homeoprotein blocks degeneration in adult dopaminergic neurons through LINE-1 repression. Embo J. 37 (2018), doi:10.15252/embj.201797374.

12. A. Benito-Gonzalez, F. J. Alvarez, Renshaw cells and Ia inhibitory interneurons are generated at different times from p1 progenitors and differentiate shortly after exiting the cell cycle. The Journal of neuroscience: the official journal of the Society for Neuroscience. 32, 1156–1170 (2012).

13. U. N. Ramírez-Jarquín, R. Lazo-Gómez, L. B. Tovar-y-Romo, R. Tapia, Spinal inhibitory circuits and their role in motor neuron degeneration. Neuropharmacology. 82, 101–107 (2014).

14. S. Layalle, M. Volovitch, B. Mugat, N. Bonneaud, M. L. Parmentier, A. Prochiantz, A. Joliot, F. Maschat, Engrailed homeoprotein acts as a signaling molecule in the developing fly. Development. 138, 2315–2323 (2011).

15. L. H. Comley, J. Nijssen, J. Frost-Nylen, E. Hedlund, Cross-disease comparison of amyotrophic lateral sclerosis and spinal muscular atrophy reveals conservation of selective vulnerability but differential neuromuscular junction pathology. Journal of Comparative Neurology. 524, 1424–1442 (2016).

16. R. A. Powis, T. H. Gillingwater, Selective loss of alpha motor neurons with sparing of gamma motor neurons and spinal cord cholinergic neurons in a mouse model of spinal muscular atrophy. Journal of Anatomy. 228, 443–451 (2016).

17. R. T. Ibad, J. Rheey, S. Mrejen, V. Forster, S. Picaud, A. Prochiantz, K. L. Moya, Otx2 promotes the survival of damaged adult retinal ganglion cells and protects against excitotoxic loss of visual acuity in vivo. The Journal of neuroscience: the official journal of the Society for Neuroscience. 31, 5495–5503 (2011).

18. I. L. Roux, A. H. Joliot, E. Bloch-Gallego, A. Prochiantz, M. Volovitch, Neurotrophic activity of the Antennapedia homeodomain depends on its specific DNA-binding properties. Proceedings of the National Academy of Sciences. 90, 9120–9124 (1993).

19. U. Bandyopadhyay, J. Cotney, M. Nagy, S. Oh, J. Leng, M. Mahajan, S. Mane, W. A. Fenton, J. P. Noonan, A. L. Horwich, RNA-Seq Profiling of Spinal Cord Motor Neurons from a Presymptomatic SOD1 ALS Mouse. Plos One. 8, e53575 (2013).

20. R. Yilmaz, K. Müller, D. Brenner, A. E. Volk, G. Borck, A. Hermann, T. Meitinger, T. M. Strom, K. M. Danzer, A. C. Ludolph, P. M. Andersen, J. H. Weishaupt, T. G. A. network MND-NET, SQSTM1/p62 variants in 486 familial ALS patients from Germany and Sweden. Neurobiol Aging (2019), doi:10.1016/j.neurobiolaging.2019.10.018.

21. M. Beurdeley, J. Spatazza, H. H. C. Lee, S. Sugiyama, C. Bernard, A. A. D. Nardo, T. K. Hensch, A. Prochiantz, Otx2 Binding to Perineuronal Nets Persistently Regulates Plasticity in the Mature Visual Cortex. Journal of Neuroscience. 32, 9429–9437 (2012).

22. U. Nordström, G. Beauvais, A. Ghosh, B. C. P. Sasidharan, M. Lundblad, J. Fuchs, R. L. Joshi, J. W. Lipton, A. Roholt, S. Medicetty, T. N. Feinstein, J. A. Steiner, M. L. E. Galvis, A. Prochiantz, P. Brundin, Progressive nigrostriatal terminal dysfunction and degeneration in the engrailed1 heterozygous mouse model of Parkinson's disease. Neurobiology of Disease. 73, 70–82 (2015).

23. M. Hossaini, S. C. Cano, V. van Dis, E. D. Haasdijk, C. C. Hoogenraad, J. C. Holstege, D. Jaarsma, Spinal Inhibitory Interneuron Pathology Follows Motor Neuron Degeneration Independent of Glial Mutant Superoxide Dismutase 1 Expression in SOD1-ALS Mice. Journal of Neuropathology and Experimental Neurology. 70, 662–677 (2011).

24. Q. Chang, L. J. Martin, Glycinergic innervation of motoneurons is deficient in amyotrophic lateral sclerosis mice: a quantitative confocal analysis. The American Journal of Pathology. 174, 574–585 (2009).

25. K. Qian, H. Huang, A. Peterson, B. Hu, N. J. Maragakis, G. Ming, H. Chen, S.-C. Zhang, Sporadic ALS Astrocytes Induce Neuronal Degeneration In Vivo. Stem Cell Reports. 8, 843–855 (2017).

26. S. C. Barber, R. J. Mead, P. J. Shaw, Oxidative stress in ALS: A mechanism of neurodegeneration and a therapeutic target. Biochimica Et Biophysica Acta Bba - Mol Basis Dis. 1762, 1051–1067 (2006).

27. O. Stettler, R. L. Joshi, A. Wizenmann, J. Reingruber, D. Holcman, C. Bouillot, F. Castagner, A. Prochiantz, K. L. Moya, Engrailed homeoprotein recruits the adenosine A1 receptor to potentiate ephrin A5 function in retinal growth cones. Development. 139, 215–224 (2012).

28. C. Kumsta, J. T. Chang, R. Lee, E. P. Tan, Y. Yang, R. Loureiro, E. H. Choy, S. H. Y. Lim, I. Saez, A. Springhorn, T. Hoppe, D. Vilchez, M. Hansen, The autophagy receptor p62/SQST-1 promotes proteostasis and longevity in C. elegans by inducing autophagy. Nat Commun. 10, 5648 (2019).

